# Working memory relates to individual differences in speech category learning: Insights from computational modeling and pupillometry

**DOI:** 10.1101/2021.01.10.426093

**Authors:** Jacie R. McHaney, Rachel Tessmer, Casey L. Roark, Bharath Chandrasekaran

**Affiliations:** Department of Communication Science and Disorders, University of Pittsburgh; Department of Speech, Language, and Hearing Sciences, University of Texas at Austin; Center for the Neural Basis of Cognition, Pittsburgh, PA

## Abstract

Across two experiments, we examine the relationship between individual differences in working memory (WM) and the acquisition of non-native speech categories in adulthood. While WM is associated with individual differences in a variety of learning tasks, successful acquisition of speech categories is argued to be contingent on *WM-independent* procedural-learning mechanisms. Thus, the role of WM in speech category learning is unclear. In Experiment 1, we show that individuals with higher WM acquire non-native speech categories faster and to a greater extent than those with lower WM. In Experiment 2, we replicate these results and show that individuals with higher WM use more optimal, procedural-based learning strategies and demonstrate more distinct speech-evoked pupillary responses for correct relative to incorrect trials. We propose that higher WM may allow for greater stimulus-related attention, resulting in more robust representations and optimal learning strategies. We discuss implications for neurobiological models of speech category learning.

## Introduction

Adults are less sensitive to speech sound contrasts that do not exist in their native language (Flege, 1995; Holt & Lotto, 2008; Iverson et al., 2003; Vallabha et al., 2007). While adults can acquire non-native speech sound categories with training (Flege, 1995; Holt & Lotto, 2006, 2010; Iverson et al., 2003; Vallabha et al., 2007; Wang et al., 1999; Wong & Perrachione, 2007), there is a great deal of individual variability in successful acquisition of speech categories in adulthood (Chandrasekaran et al., 2015), which may be driven in part by individual differences in working memory (WM) and executive attention (EA; Kidd et al., 2018). Operationally, WM refers to the ability to represent and manipulate information in a buffer that is consciously accessible (Engle & Kane, 2004; Tsukahara et al., 2016). EA provides the scaffolding for this operation by enabling the maintenance of information and preventing interference from external sources (Engle & Kane, 2004). Tasks like the operation span (OSPAN; Unsworth et al., 2005) can be used to measure both WM and EA, but we will refer to this measure solely as WM henceforth. Within a population, there is consistent variation in WM that systematically relates to the successful acquisition of various components of language (Kidd et al., 2018). However, the precise role of WM in mediating individual differences in speech category acquisition has not been established. Across two experiments, we examine the extent to which individual differences in WM relate to successful acquisition of a non-native speech contrast (Mandarin tones). The first experiment is a large-scale experiment conducted online (*n* = 196 participants) where we examined the role of WM in speech category learning. In a second experiment conducted in-person (*n* = 28), we leveraged pupillometry and computational modeling to specify mechanisms and computational strategies driving a potential WM advantage in speech category learning success.

Despite the individual differences in learning success, most adults can acquire even perceptually subtle non-native contrasts with training. Explicit training regimens that have engendered significant, generalizable learning, typically incorporate three characteristics: natural speech productions, talker and contextual-variability, and feedback (Bradlow, 2008). The use of natural speech productions and multiple talkers and contexts are beneficial because they expose learners to the natural variability in the learning environment, allowing learners to focus on and weigh salient dimensions that vary less across talkers (Bradlow, 2008; Bradlow & Bent, 2008). Listeners may use feedback to monitor errors, fine-tune their learning strategies to focus on relevant information, and/or ignore irrelevant information. However, the extent and quality of feedback required is unclear. While adults require some amount of feedback to learn (Chandrasekaran et al., 2015; Chandrasekaran, Yi, et al., 2014; Lim & Holt, 2011; McClelland et al., 2002; Tricomi et al., 2006; Yi et al., 2016), adults can acquire speech categories incidentally (Y. Gabay et al., 2015; Lim et al., 2019; Luthra et al., 2019) or with task-irrelevant feedback (Goudbeek et al., 2008; McClelland et al., 2002).

In explicit training regimens, individual differences in speech category learning success could arise from individual variation in the ability to select relevant dimensions and ignore irrelevant or distracting dimensions. Individuals with lower WM demonstrate a relatively reduced ability to attend to and select task-appropriate dimensions (D’Esposito & Postle, 2015; Unsworth & Robison, 2017b; Wöstmann & Obleser, 2016). A recent framework posits that individual differences in WM are driven by moment-to-moment fluctuations in locus coeruleus-norepinephrine (LC-NE) system activity (Unsworth & Robison, 2017a). Individuals with lower WM may have greater attentional fluctuations that are likely caused by a disruption in LC-NE functioning when compared to individuals with higher WM (Unsworth & Robison, 2017a). The LC-NE system receives reciprocal input from the prefrontal cortex, such that these reciprocal projections can modulate attention for salient and irrelevant events (Aston-Jones & Cohen, 2005; Berridge & Waterhouse, 2003). Thus, individuals with lower WM may show reduced attention to salient dimensions relative to those with higher WM, resulting in less effective weighting of task-relevant dimensions during category learning.

In addition to attention to relevant dimensions, individual differences in speech category learning success in explicit training regimens could also arise from individual variation in the use of corrective feedback to guide appropriate learning strategies. Individual differences in WM may impact the balance of learning strategies during training. Adapted from a model of multiple learning systems in the visual domain (Ashby et al., 1998), the dual-learning systems (DLS) framework posits that at least two competitive learning systems mediate feedback-based learning of novel speech categories in adulthood (Chandrasekaran, Koslov, et al., 2014; Chandrasekaran, Yi, et al., 2014; Feng, Yi, et al., 2018; Maddox & Chandrasekaran, 2014). A *reflective* learning system, involving the prefrontal cortex and the hippocampus, uses WM (Decaro et al., 2008; Filoteo et al., 2010; Miles et al., 2014; Reetzke et al., 2016; Zeithamova & Maddox, 2006) to develop and test verbalizable rules based on feedback (Maddox & Ashby, 2004). In contrast, a *reflexive* learning system that is independent of WM, associates perception with rewarding actions that are derived from feedback (Maddox & Chandrasekaran, 2014; Seger, 2008; Seger & Miller, 2010). The reflexive learning system requires pre-decisional integration across perceptual dimensions for optimal categorization (Ashby & Gott, 1988; Ashby & Maddox, 2011).

The reflective and reflexive learning systems are argued to be competitive throughout learning, though there is a slight bias for the reflective system during early learning in humans (Chandrasekaran, Koslov, et al., 2014; Maddox & Chandrasekaran, 2014). Learners often begin by using the reflective learning system and rule-based learning strategies for categorization but switch to the reflexive learning system if the categories are more optimally acquired via procedural-based learning strategies (Ashby et al., 1998). Prior studies have leveraged computational models to show that during the acquisition of non-native speech categories, successful learners tend to switch to procedural-based learning strategies, while less successful learners persevere with suboptimal rule-based learning strategies (Chandrasekaran, Koslov, et al., 2014; Maddox & Chandrasekaran, 2014). Individuals with higher WM may be less likely to use suboptimal rule-based learning strategies, allowing for a faster transfer of control to the reflexive learning system. An alternate possibility, derived from the perspective that the two learning systems are competitive, is that individuals with lower WM may switch to procedural-based learning strategies, which are independent of WM, and demonstrate an *advantage* in learning nonnative speech strategies. Indeed, the role of WM during procedural-based category learning remains unresolved. While some studies have shown that procedural-based learning is impaired by increasing WM demands (Miles et al., 2014; Zeithamova & Maddox, 2006; but see Newell et al., 2010), others have demonstrated that the procedural-based learning system is *facilitated by* WM demands (Filoteo et al., 2010; but see Newell et al., 2013). Based on this conflicting evidence, the extent to which WM influences feedback-based speech category learning strategies is unclear and warrants systematic investigation.

The current study utilizes a combination of behavioral, physiological, and computational approaches to examine the extent to which individual differences in WM are associated with the acquisition of Mandarin tone categories by native English speakers. In Mandarin, pitch changes within a syllable can differentiate word meaning. Mandarin has four linguistically-relevant pitch contrasts, or tones, that function similarly to consonants and vowels in modifying word meaning. At least two pitch dimensions, relative pitch and pitch change, phonetically contrast the tone categories (Chandrasekaran et al., 2007; Gandour & Harshman, 1978). Adult English listeners often have difficulty learning Mandarin tone contrasts because these contrasts do not differentiate word meaning in English (Wang et al., 1999, 2003). Prior studies have demonstrated that neural representations of tone categories can emerge within a single session of sound-to-category training (Feng, Yi, et al., 2018; Yi et al., 2016). Specifically, Feng et al. (2018) showed rapid emergent representations of tone categories in the left superior temporal gyrus within a few hundred training trials. Interestingly, during the second half of training, there was greater functional coupling between the left superior temporal gyrus and the striatum during incorrect responses, suggesting that representations are ‘fine-tuned’ as a function of corrective feedback. Successful learners tend to show greater striatal activation as well as more robust, multidimensional category representations in the left superior temporal gyrus (Feng, Yi, et al., 2018).

In the first experiment, we predicted that WM differences in a large, highly diverse population of non-tonal language speakers would relate to individual differences in the successful acquisition of tone categories. In the second experiment, we replicate Experiment 1 in a smaller cohort but with the addition of computational modeling and pupillometry to provide mechanistic insight into the role of WM in non-native speech category learning. Computational modeling approaches allowed us to go beyond accuracies and discern underlying neurobiological learning strategies. Pupillary responses serve as an indirect index of LC-NE activity (Alnæs et al., 2014; Aston-Jones & Cohen, 2005; S. Gabay et al., 2011; Murphy et al., 2014). Prior studies have shown distinct pupillary responses as a function of individual differences in WM. Pupillary measures during the process of speech category learning allowed us to discern the extent to which LC-NE activity differences between individuals with higher and lower WM underlies variability in speech category learning success (Heitz & Engle, 2007; Tsukahara et al., 2016).

In both experiments, we predict that individuals with higher WM would show greater tone category learning, and greater generalizability to untrained talkers relative to those with lower WM. In Experiment 2, we predict that individuals with higher WM would show a faster switch to optimal procedural-based learning strategies, and that this change would be discerned online via pupillometric changes. We also predicted that the reflective learning system scaffolds the reflexive learning system, thus allowing individuals with higher WM to switch faster from suboptimal rule-based learning strategies to optimal procedural-based learning strategies. In line with the LC-NE hypotheses, we predict distinct stimulus-evoked pupillary differences between individuals with higher and lower WM. We predict that those with lower WM would show greater moment-to-moment attentional fluctuations during sound-to-category learning. Pupillary responses tend to be larger for incorrect trials than for correct trials during behavioral tasks (Braem et al., 2015; Critchley et al., 2005), therefore, we predict less distinct stimulus-evoked pupillary responses during correct trials in individuals with lower WM compared to incorrect trials. We also predict that a greater shift to optimal procedural-based learning strategies in individuals with higher WM manifests as a larger training-related reduction in pupil dilation, reflecting the reduced cognitive effort associated with procedural-based learning strategies. Overall, we sought to provide novel mechanistic insights into an important variable (i.e., WM) that may underlie individual differences in speech category learning success.

## Experiment 1

### Methods

#### Participants

A total of 198 participants between the ages of 18-35 (99 females; mean age = 24.97, *SD* = 4.97) were recruited online using Prolific (www.prolific.co). Prolific provides access to participants from all over the world for a representative study sample. Participants first completed a language-history questionnaire to ensure they reported no prior experience with a tonal language (e.g., language courses, immersion experiences). Participants received monetary compensation for their participation. Two participants did not follow task instructions and were removed from data analysis, resulting in a final sample of 196 participants (98 females; mean age = 24.939, *SD* = 4.934). This research protocol was approved by the Institutional Review Board at the University of Pittsburgh.

#### Stimuli

Participants learned to categorize Mandarin tones that vary along relative pitch and pitch change dimensions. Four lexical tones were produced in five syllable contexts (/bu/, /di/, /lu/, /ma/, /mi/) by four native Mandarin Chinese speakers (two females; Fig. 1A; Feng, Gan, et al., 2018). The Mandarin tone stimuli were RMS amplitude (70 dB) and duration (440 ms) normalized using Praat (Boersma & Weenink, 2005). A scatterplot of the speech stimuli is displayed in Figure 1B.

**Figure 1.**
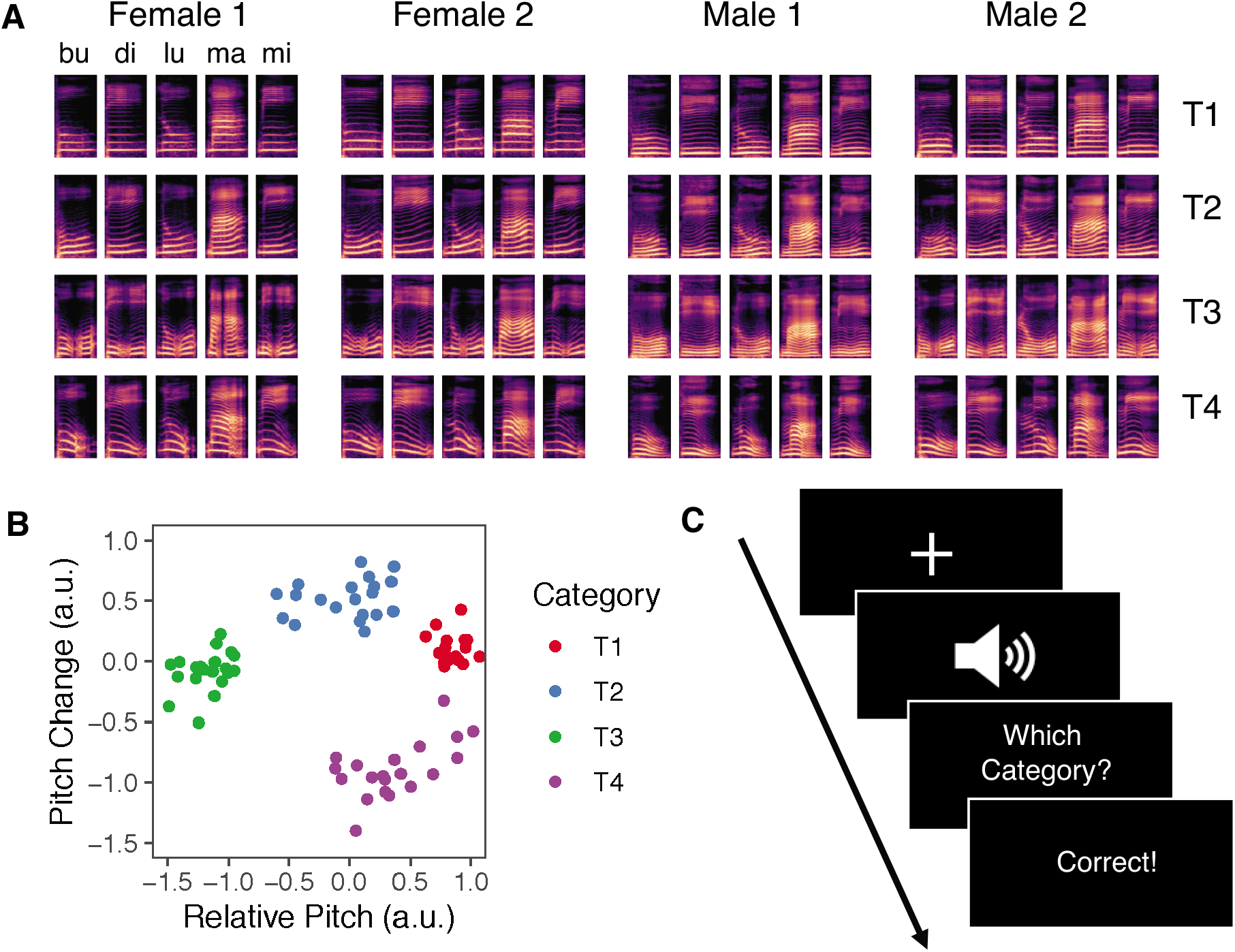
Experimental stimuli and training procedure. A) Spectrograms for four Mandarin tones (T1, T2, T3, T4) produced by two female and two male native speakers in five syllable contexts: /bu/, /di/, /lu/, /ma/, and /mi/. B) Scatterplot of all stimuli from the training and generalization blocks along the two acoustic dimensions of relative pitch and pitch change in arbitrary units (a.u.). C) Representative trial of the Mandarin tone training task procedure.

#### Procedure

The experiment was created and hosted using Gorilla Experiment Builder (Anwyl-Irvine et al., 2020). Participants were instructed to use headphones for the training and generalization tasks and were instructed to set the stimulus volume to a comfortable listening level. Each trial in the training task began with a fixation cross (1000 ms), followed by the successive presentation of the auditory stimulus (440 ms), category response (untimed), and corrective feedback (750 ms; Fig. 1C). Feedback informed participants whether their response was “Correct” or “Wrong.” The training task consisted of six blocks of 10 trials of each of the four tone categories produced by two native Mandarin speakers (one female). Each block contained 40 trials for a total of 240 trials across the training task. Participants received a self-timed break at the end of each block. A generalization task immediately followed the training task. The generalization task consisted of a single block of 10 trials of each of the four tone categories (40 trials total) with no corrective feedback and used speech stimuli that were produced by two novel native Mandarin speakers (one female) that were not used in the training blocks.

All participants also completed the automated version of the operation span (OSPAN) task (Unsworth et al., 2005). The OSPAN task has been widely used to measure WM and EA (e.g., Turner & Engle, 1989; Xie et al., 2015). Participants were shown simple arithmetic problems and asked to decide whether presented solutions to the problems were correct or incorrect. A letter was displayed on the screen after each problem. Following a series of arithmetic problems, participants were required to recall the letters that were displayed in the order that they appeared. The task consisted of 15 letter sequences that spanned three to seven letters (three repetitions of each span). If a participant correctly recalled all letters from a sequence, the span length was added to their score. For instance, if a participant recalled all letters in the correct order from a seven-letter span sequence, seven points were added to their score. The maximum possible score on the OSPAN task was 75. Participants were then divided into higher (n = 101) or lower (n = 95) WM groups based on a median split (median OSPAN score= 45; Fig. 2A; Zinke, Zeintl, Eschen, Herzog, & Kliegel, 2012). OSPAN scores were also analyzed as a continuous measure in the behavioral data analysis to ensure that the median split accurately captured individual differences in WM.

**Figure 2.**
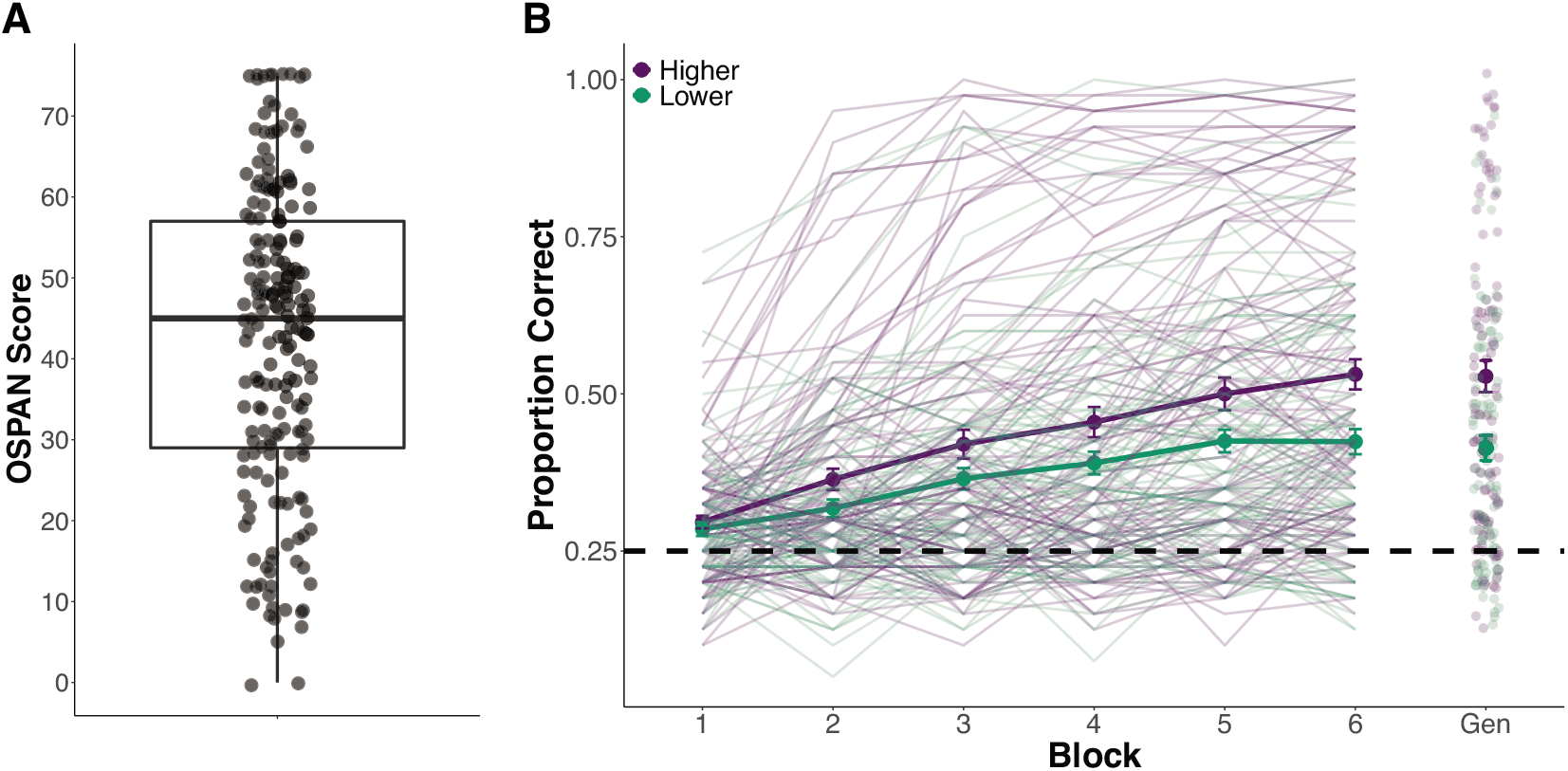
Experiment 1 behavioral results. A) Distribution of scores on the OSPAN task for all 196 participants. B) Categorization accuracy across the six category training blocks and the generalization block (abbreviated as “Gen”) for higher and lower working memory (WM) participants. The dark, solid lines denote the average accuracy with error bars reflecting standard error of the mean for each WM group. The lighter lines and points represent each individual participant’s accuracies. Categorization accuracy is defined as the proportion of correct trials per block. The dashed line reflects chance-level performance (0.25).

#### Behavioral data analysis

A binomial generalized linear mixed-effects model was fit to examine learning performance in the tone categorization training task using the lme4 package (Bates et al., 2015) in R (R Core Team, 2019) and *p*-values were estimated using the lmerTest package (Kuznetsova et al., 2017). The outcome variable was trial-by-trial accuracy (correct, incorrect) for each participant. The optimal model fit included fixed effects of trial, WM (reference = lower WM), and the interaction of trial and WM, with random intercepts of subject and tone category.

We fit a similar binomial generalized linear mixed-effects model to examine categorization accuracy in the generalization block. However, this model varied in that the outcome variable was trial-by-block response outcomes in the generalization block and block 1 of the training task. Block 1 was used as the reference block to account for individual differences in performance at baseline and to rule out any differences in categorization accuracy between higher and lower WM participants at the onset of training. The model included fixed effects of block, WM, and the interaction of block and WM, with random intercepts of subject and tone category.

### Results

Participants with higher WM had greater overall accuracy in the category learning training task (β = 0.272, *z* = 2.467, *p* = 0.014) relative to participants with lower WM. Additionally, there was a positive and significant effect of trial for the participants with higher WM (β = 0.142, *z* = 6.893, *p* < 0.001). These results suggest that, not only did the participants with higher WM demonstrate greater categorization accuracy throughout the training task, but they also had a larger trial-by-trial improvement in categorization accuracy relative to participants with lower WM (Fig. 2B). When examining WM as a continuous variable, we found that individual participants’ OSPAN scores significantly correlated with categorization accuracy in block 4 (*r* = 0.190, *p* = 0.009), block 5 (*r* = 0.200, *p* = 0.005), and block 6 (*r* = 0.230, *p* = 0.001) of training, but did not significantly correlate with categorization accuracy in early training blocks (*p*s > 0.05). These significant correlational results support the findings based on the WM median split and demonstrate that participants with higher WM learned non-native speech categories to a greater degree than participants with lower WM.

We also examined differences in categorization accuracy between higher and lower WM participants in the generalization block, wherein participants categorized stimuli produced by novel speakers without corrective feedback. There was a positive and significant interaction of block and higher WM (β = 0.472, *z* = 6.671, *p* < 0.001). This result suggests that the participants with higher WM had a greater improvement in categorization accuracy from baseline to the generalization block than the participants with lower WM. Additionally, there were no differences in categorization accuracy between higher and lower WM participants in block 1 of the training task (β = 0.027, *z* = 0.272, *p* = 0.786), demonstrating that differences in improvement by the generalization block cannot be attributed to differences in categorization accuracy between these groups at the onset of training. Moreover, categorization accuracy in the generalization block also positively correlated with individual participants’ OSPAN scores (*r* = 0.260, *p* < 0.001). This significant correlation suggests that individuals with higher WM were better able to generalize learned categories to a context involving untrained talkers.

## Experiment 2

Experiment 1 demonstrated significant differences in non-native speech categorization performance based on WM status. In Experiment 2 we probe the mechanisms underlying the WM advantage to learning tone categories. Experiment 2 takes a computational and mechanistic approach to examine the role of the LC-NE system, as indexed by task-evoked pupillary responses, on WM involvement during non-native speech category learning.

### Methods

#### Participants

Twenty-eight participants (21 females; mean age = 20.54, *SD* = 3.00) were recruited from The University of Texas at Austin and the greater Austin community for Experiment 2. All participants were native English listeners, had hearing thresholds <25 dB for 250, 500, 1000, 2000, 4000, and 8000 Hz, and had less than six years of formal music training. Participants first completed a language-history questionnaire to ensure no prior tonal language experience (e.g., language courses, immersion experiences). Participants received either monetary compensation or research credit for their participation. This research protocol was approved by the Institutional Review Board at The University of Texas at Austin.

#### Procedure

The speech stimuli for the training task were identical to those used in Experiment 1 (see Experiment 1 *Stimuli* section). In addition to the four Mandarin tone categories, we also included a fifth category of silent trials to examine the extent to which the observed pupillary responses were sound-evoked and not caused by other physiological responses (e.g., motor preparation) This increased the number of trials per block to 50 (40 speech trials, 10 silence trials), for a total of 300 trials across six blocks of training. Mapping of the category and key response was counterbalanced across participants, with a different category designated as a no-press response. For instance, some participants were instructed to not press a key if they believed the stimulus belonged in category 1, but to press keys ‘2’, ‘3’, and ‘4’ for categories 2, 3, and 4, respectively. Key 5 was designated for silence trials. The training task was created and presented using Experiment Builder (SR Research), and the speech stimuli were presented via insert earphones (ER-3C; Etymotic Research, Elk Grove Village, IL).

Monocular left eye pupil size was monitored using an Eyelink 1000 Plus Desktop Mount with a chin and forehead rest for stabilization. Data were recorded at a sampling rate of 1000 Hz. Luminance of the visual field was controlled by setting consistent room lighting across all participants. Nine-point eye tracker calibration was performed prior to the start of the experiment. Consistent with procedures from Experiment 1, each trial began with a fixation cross, followed by the speech stimulus presentation, category response, and corrective feedback. However, we included delays after each trial event to allow for changes in pupil size (Koelewijn et al., 2018; Winn et al., 2018; Zekveld et al., 2013). Participants were required to fixate on the cross in the center of the screen for a minimum of 500 ms to begin each trial in an effort to minimize pupil foreshortening errors (Hayes & Petrov, 2016). The auditory stimulus was presented following a two-second delay after meeting the fixation criteria. There was a four-second delay from the onset of the auditory stimulus to the category response prompt on the screen in the form of: “Which Category?”. Participants had two seconds to respond. Following the response, there was a two-second delay before corrective feedback was displayed on the screen in the form of “Correct” or “Wrong” for two seconds. At the end of each block, tone categorization accuracy from the previous block was displayed onscreen in the form of a percentage. Drift correction was performed between each block. Following the training task, participants completed a tone categorization generalization block and the OSPAN task as described in Experiment 1 *Procedure* section. Analysis of the behavioral data from the category learning task was identical to Experiment 1 (see Experiment 1 *Behavioral data analysis* section), with the exception that silence trials were removed for behavioral analyses. Similar to Experiment 1, participants were then categorized into higher (*n* = 14) and lower (*n* = 14) WM groups based on a median split (median OSPAN score = 41). OSPAN scores were additionally assessed as a continuous measure as in Experiment 1.

Upon completion of the training task and the generalization block, participants filled out the NASA-task load index (NASA-TLX; Hart & Staveland, 1988) questionnaire. Participants completed it as a self-report measure to assess effort, frustration level, mental demand, physical demand, temporal demand, and overall performance on the speech category learning task (Table 1).

**Table 1.** Experiment 2 Demographics

| Demographic | Higher<br>Working Memory | Lower<br>Working Memory | *t* | *p* |
| --- | --- | --- | --- | --- |
| *n* (Female) | 14 (8) | 14 (13) | - | - |
| Age (Mean, SD) | 21.500 (3.94) | 19.571 (1.089) | 1.766 | .098 |
| OSPAN (Mean, SD) | 52.071 (7.022) | 27.286 (8.453) | 8.440 | < .001 *** |
| NASA-TLX (Mean, SD) |  |  |  |  |
| Effort | 73.571 (11.507) | 68.571 (19.158) | 0.837 | .412 |
| Frustration Level | 58.571 (22.311) | 47.857 (28.333) | 1.112 | .277 |
| Mental Demand | 67.143 (24.629) | 74.643 (12.929) | -1.009 | .325 |
| Overall Performance | 46.786 (26.502) | 47.143 (22.678) | -0.038 | .970 |
| Physical Demand | 41.071 (21.766) | 33.214 (23.420) | 0.919 | .366 |
| Temporal Demand | 30.000 (19.315) | 41.071 (22.203) | -1.408 | .171 |

\*\*\* $p < .001$

#### Decision Bound Computational Modeling

While behavioral accuracies provide important insights, they provide no information about *how* participants learned the categories and what strategies they used. To understand the strategies participants used, we applied several classes of decision-bound computational models (Ashby, 1992; Ashby & Maddox, 1993; Chandrasekaran et al., 2016; Smayda et al., 2015). Decision bound models assume that participants divide the categories in the two-dimensional stimulus space (i.e., relative pitch and pitch change) with decision boundaries that can rely on rule-based or procedural-based learning processes.

Originally developed in the visual modality, decision bound models were recently extended to examine auditory category learning (Chandrasekaran et al., 2016; Goudbeek et al., 2008, 2009; Maddox et al., 2002, 2013, 2014; Maddox & Chandrasekaran, 2014; Roark & Holt, 2019; Scharinger et al., 2013; Smayda et al., 2015). These models are fit individually for each participant and each block to mitigate challenges interpreting fits to aggregate data (Ashby et al., 1994; Estes, 1956; Estes & Maddox, 2005). Consistent with prior research, we specified three classes of models, with multiple instantiations possible within a class (Chandrasekaran et al., 2015, 2016; Maddox et al., 2014, 2016; Smayda et al., 2015; Yi et al., 2016). The classes included *Rule-Based models, the Striatal Pattern Classifier model* (*SPC*), and the *Inconsistent/Random* Responder models.

##### Rule-Based Models

The Rule-Based class of models represents hypothesis-testing mechanisms that are dependent on WM resources and mediated by the reflective learning system. Rule-Based models assume that participants selectively attend to individual dimensions during learning and can be either unidimensional (attending to a single dimension) or multidimensional (attending to both dimensions).

Unidimensional models assume that participants draw boundaries between categories based only on one of the dimensions (i.e., relative pitch or pitch change). Unidimensional models have four free parameters: three reflect the placement of the decision boundaries along the relevant dimension and one reflects perceptual and criterial noise. Separate unidimensional models were fit assuming participants made categorization decisions using only the relative pitch dimension or only the pitch change dimension.

Multidimensional models assume that the participant places two decision boundaries (one along each dimension) that are combined to determine category membership. When a participant uses a multidimensional strategy, they selectively attend to both dimensions. One subclass of multidimensional models has three free parameters: two reflect the placement of a single boundary along each dimension and one reflects perceptual and criterial noise. Models from this subclass are also referred to as ‘conjunctive’ models as the boundaries require the conjunctive combination of the boundaries along the two dimensions to separate all four categories and forms the shape of a plus sign (Fig. 3). A separate subclass of multidimensional models has four free parameters: two reflect the placement of two boundaries along one dimension, one reflects the placement of a single boundary along the other dimension, and one reflects perceptual and criterial noise. Models from this subclass are sometimes called ‘conjunctive-H’ models as the boundaries form the shape of an H (Fig. 3).

**Figure 3.**
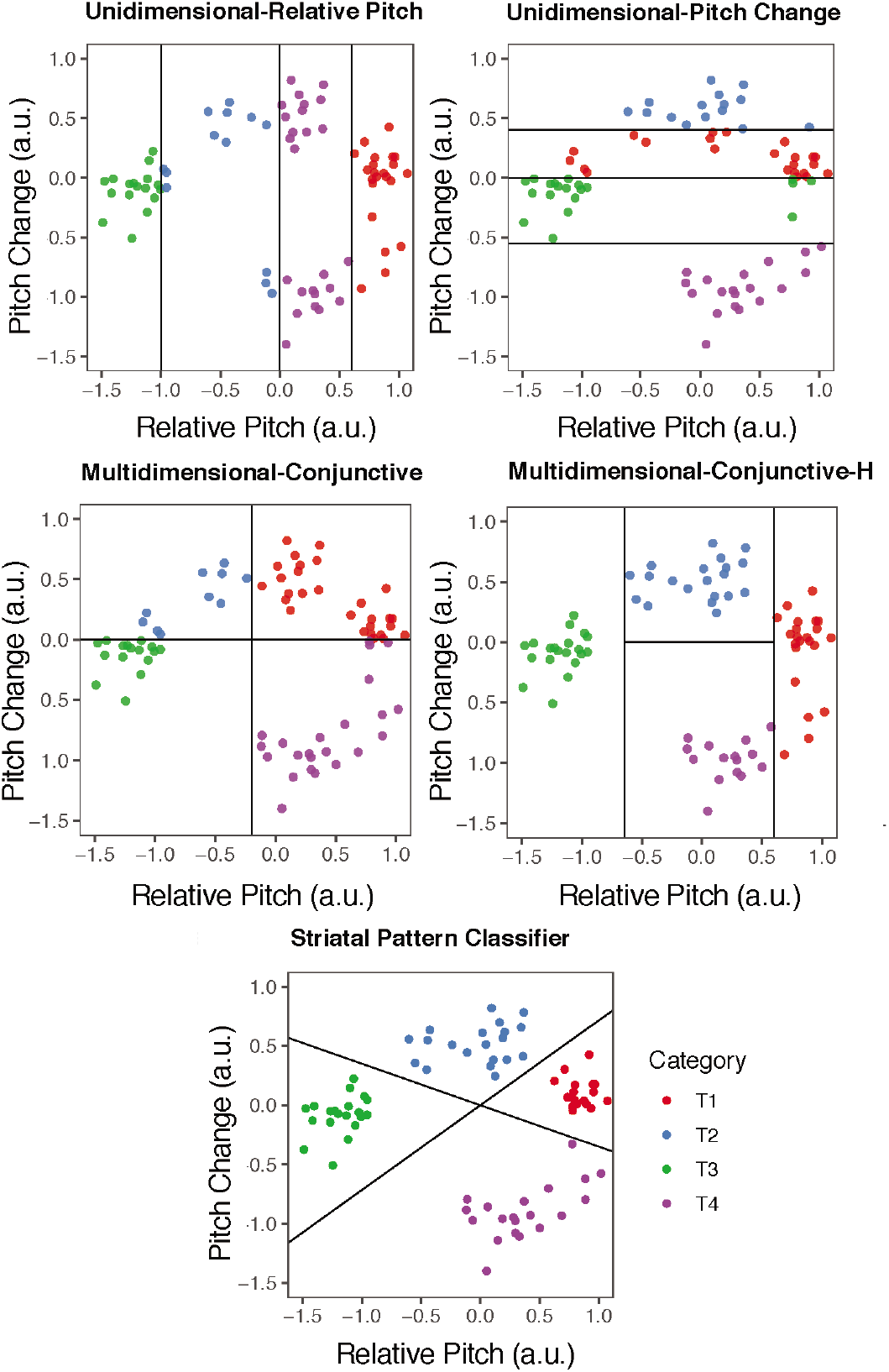
Hypothetical responses to each stimulus in the Mandarin tone category space (T1, T2, T3, T4), where each colored dot is a stimulus and the color corresponds to the theoretical response of a participant. Each panel demonstrates how the models make sense of the participant response data by placing decision boundaries in the space based on the participant’s responses. This will be more or less accurate depending on the participant’s actual response patterns and the model that is being fit. In each plot above, the stimuli are the same; it is only the mapping of stimulus to response label that is different across the panels. This is how different participants are best-fit by different models and we can conclude that they are using different strategies to separate the categories.

##### Striatal Pattern Classifier Model

The implicit Striatal Pattern Classifier (SPC) model is a neurobiologically grounded model thought to represent procedural-based learning mechanisms (Ashby et al., 1998). The SPC model assumes that participants use feedback to learn stimulus-response associations instantiated within the striatum (Ashby & Waldron, 1999). The SPC model assumes that participants build category representations based on theoretical ‘striatal’ units (based on the neurobiology of medium-spiny neurons), where similar category responses to stimuli are clustered together in perceptual space, supported by the reflexive learning system (Ashby & Waldron, 1999). The SPC model assumes participants use both dimensions to divide the categories and that the boundaries are not orthogonal to the dimensions (Fig. 3).

The SPC model has nine free parameters: eight that determine the location of theoretical ‘striatal’ units in perceptual space (x and y coordinates) and one that represents the noise associated with the placement of the striatal units. When a participant uses an SPC strategy, they combine information across both dimensions in a pre-decisional manner. The SPC model is considered the optimal model for learning Mandarin tone categories (Yi et al., 2016) and has been linked to a corticostriatal pathway that supports learning (Ashby & Ell, 2001; Nomura & Reber, 2008; Yi et al., 2016).

##### Inconsistent/Random Responder Model

The Inconsistent or Random Responder model assumes that participants respond to the stimuli with all responses being equally probable. Thus, this model captures participants who respond randomly to the stimuli and participants who may be changing strategies too often to be accounted for by the other models. We refer to these participants as Inconsistent strategy users.

##### Summary

The different classes of decision bound models make different assumptions about how participants divide the stimuli into categories based on the underlying dimensions that define the stimuli. As a demonstration of these models, Figure 3 shows a hypothetical response pattern for each model class. Mathematical formulation of these models is outside of the scope of the current article and can be found elsewhere (Ashby, 1992; Ashby & Maddox, 1993).

As input, each model uses the dimensional coordinates (i.e., relative pitch and pitch change) of each stimulus and the participant’s actual response to that stimulus for a given block of each participant’s data. Importantly, these models are agnostic about the actual category identity of the stimuli and are based on the participant’s response. To visualize this, for each block trials, we can plot all of the stimuli that the participant encountered in the two-dimensional space and divide that space using decision boundaries (Fig. 3) with each of the classes of models that have different assumptions with the goal of finding the best model to account for this particular participant’s response pattern in this block of data. The key differences between the models are in the assumptions they make about how participants use the stimulus dimensions to separate the categories. In some instances, they use only one dimension (unidimensional rule-based models). In other cases, they use both dimensions in a manner that is orthogonal to the dimension itself, consistent with selectively attending to that dimension (multidimensional rule-based models). Finally, in the SPC case, they use both dimensions in a manner that is not orthogonal to the dimensions, but rather is a relationship that is harder to describe verbally and requires integration across the dimensions.

##### Model Fitting and Selection

For each of the classes of models described above, several versions were fit that made different assumptions about assignment of responses to regions of space. In total, 11 unidimensional models, 13 multidimensional rule-based models, one SPC model, and one random responder model were fit to each block of each participant’s data. Models were fit to the early and late blocks for each participant. As such, this leads to many models being fit across participants (26 models x 28 participants x 2 blocks = 1,456 models fit).

Model parameters were estimated using maximum likelihood procedures (Wickens, 1982) and model selection used the Bayesian Information Criterion (BIC), which penalizes models with more free parameters: BIC = *r*\*ln*N* - 2ln*L*, where *r* is the number of free parameters, *N* is the number of trials in a given block for a given subject, and *L* is the likelihood of the model given the data (Schwarz, 1978). Within each block for each participant, the one model among all 26 individual models from all classes with the lowest BIC value was selected as the best-fit model.

##### Assessment of Goodness of Fit

To understand how well the best-fit model for each participant and block accounted for that participant’s actual response pattern, we computed the proportion of participant’s responses accurately predicted by the best-fit model (i.e., model accuracy) as a measure of goodness-of-fit.

Once the best-fitting model was selected using BIC (comparing against all other models), we computed the *predicted* response based on the fitted parameters for the best-fit model. That is, as these models are categorical models, this is the category response (i.e., T1, T2, T3, or T4) that the best-fitting model predicts that the participant would have made if they applied this strategy consistently throughout that block. Next, we compared the predicted response of the best-fitting model to the *observed* response made by the participant for each stimulus presented in a given block. For example, for Stimulus-1, if the model predicted that the participant would have responded 1 and the participant actually responded 1, this would be coded as a ‘Correct’ prediction. However, if the model predicted 1 and the participant actually responded 2, this would be coded as an ‘Incorrect’ prediction. For each participant and each block, we computed the model accuracy by comparing the best-fit model’s predicted response and the participant’s actual (observed) response.

Using this goodness-of-fit measure of model accuracy, we observed that the best-fit models accurately captured participants’ response patterns with an average predicted vs. observed accuracy of 50% in the early block and 61% in the late block, which were substantially better than chance (25%). Because the accuracy was better than chance, we know that the best-fit models provided an accurate account of participants’ response patterns. It was expected that the model accuracy would be less than 100% (i.e., not account for every single one of the participant’s responses in a given block of data). The reason model accuracy may be less than 100% is if a participant applied the strategy even somewhat inconsistently across all trials or they had a lapse in attention during even a single trial. Instead, the approach of these models was to capture the most likely strategy across a given block of trial and provide an estimate (with relatively high accuracy) of the strategy participants were using over the block of trials.

#### Pupillometry pre-processing

Consistent with prior research, pupillometry data were preprocessed to remove noise from the analysis (Winn et al., 2015; Zekveld & Kramer, 2014). Pupillometry data were down-sampled to 50 Hz (Wierda et al., 2012) and trials with more than 15% of samples detected as blinks were removed (n = 2044 out of 8400 training trials; McMahon et al., 2016). Missing samples due to blinks were linearly interpolated to 120 ms before and after the blink. Additional blinks were identified and linearly interpolated based on the first derivative of the blink threshold. Pupil responses were baseline normalized using the average pupil size in the 500 ms prior to the onset of the speech stimuli (Peysakhovich et al., 2015), wherein the outcome variable reported is the proportion change in pupil size relative to baseline.

#### Growth curve analysis

Pupil responses within the time window from 0 to 2500 ms time-locked to the speech stimulus onset were analyzed using growth curve analysis (GCA; Mirman, 2014). GCA is appropriate for modeling time-series data, such as pupillary responses, because it provides a statistical approach for modeling changes in the timing and shape of the pupillary response over time. GCA has an advantage over traditional approaches, like time-binned analysis of variance, for several reasons: 1) GCA does not require time-binned samples, which eliminates the trade-off between temporal resolution and statistical power, 2) there is no experimenter bias in selecting time windows in an arbitrary manner, and 3) the model can account for individual differences (Mirman, 2014).

The pupillary response unfolds over time and often does not follow a linear trajectory (Winn et al., 2015, 2018). Therefore, GCA uses orthogonal polynomial time terms to capture distinct functional forms of the pupillary response. A GCA was fit to model the interactions between first-, second-, third-, and fourth-order orthogonal polynomials and the independent variables of interest. The fourth-order polynomial was chosen to model the pupillary response because it was consistent with the trajectory of the data and provided a better fit than a model that included the only first-, second-, and third-order polynomials (*χ*^2^(6) = 49.115, *p* < .001; AIC_M1_ = −21,288; AIC_M2_ = −21,325). This fourth-order model uses four parameters to capture the complexity of the pupillary response. The intercept refers to the overall change in the pupillary response over the entire time window. The linear (ot1) term reflects the slope of the pupillary response. The quadratic (ot2) term represents the curvature of the pupillary response. Given the trajectory of the pupillary response, the quadratic term should be negative. Here, a larger, negative quadratic term reflects a steeper curvature. Lastly, the cubic (ot3) and quartic (ot4) terms represent the extent to which two or three inflection points occur in the pupillary response, respectively. GCA were conducted using the lme4 package (Bates et al., 2015) with log-likelihood maximization using the BOBYQA optimizer to promote convergence, and *p*-values were estimated using the lmerTest package (Kuznetsova et al., 2017).

We first estimated a GCA to examine differences in pupillary responses between sound trials and silence trials. The optimal, final model included fixed effects of trial type (sound or silence; reference = silence) on all time terms with random slopes of subject on each time term and trial type. This random effect structure provided a better model fit than when including only random slopes of subject on each time term (*χ*^2^(6) = 5,763.000, *p* < .001; AIC_M1_ = −34,269; AIC_M2_ = - 40,020).

We modeled a second GCA to examine pupillary changes for higher WM and lower WM participants on correct and incorrect trials between the early and late halves of training. This model included fixed effects of WM (reference = lower WM), trial accuracy (reference = incorrect trials), and training half (reference = early half) on all time terms, and the interaction of WM, accuracy, and half on all time terms. This model also included random slopes of subject on each time term and accuracy type, which provided a better model fit than including only random slopes of subject on each time term (*χ*^2^(6) = 2,146.800, *p* < .001; AIC_M1_ = −53,572; AIC_M2_ = −55,707).

### Results

#### Behavioral

Consistent with the findings from Experiment 1, participants with higher WM had significantly greater overall average accuracy (β = 0.763, *z* = 2.233, *p* = 0.026) and a larger trial-by-trial improvement in accuracy (β = 0.173, *z* = 2.994, *p* = 0.003) relative to participants with lower WM during the training task (Fig. 4B). Individual participants’ OSPAN scores positively correlated with categorization accuracy in block 2 (ρ = 0.480, *p* = 0.010), block 3 (ρ = 0.420, *p* = 0.028), block 4 (ρ = 0.380, *p* = 0.044), and block 5 (ρ = 0.410, *p* = 0.030), but were not statistically significant for block 1 (ρ = 0.015, *p* = 0.940) or block 6 (ρ = 0.350, *p* = 0.071). These findings suggest that participants with higher OSPAN scores tended to have higher categorization accuracy for most of the training task. In the generalization block, there was a significant and positive interaction of accuracy and higher WM participants (β = 0.429, *z* = 2.127, *p* = 0.033), which demonstrates that the improvement in categorization accuracy from block 1 to the generalization block was greater for higher WM participants relative to lower WM participants. There was no significant effect of accuracy in block 1 between higher and lower WM participants (β = 0.042, *z* = 0.133, *p* = 0.894), suggesting that the improvement observed from block 1 to the generalization block in the higher WM participants was not primarily driven by differences in baseline performance. Collectively, the results from the categorization task in Experiment 2 are similar to the findings from Experiment 1 and demonstrate participants with higher WM have better Mandarin tone category learning than those with lower WM. In contrast to the findings from Experiment 1, individual participants’ OSPAN scores did not significantly correlate with accuracy in the generalization block (p = 0.270, *p* = 0.160) in Experiment 2. Additionally, higher and lower WM participants did not significantly differ on any of the NASA-TLX subscales (see Table 1), which indicates that both WM groups found the speech category learning task equally demanding on workload. Thus, the differences observed between higher and lower WM participants derive from specific aspects of learning.

**Figure 4.**
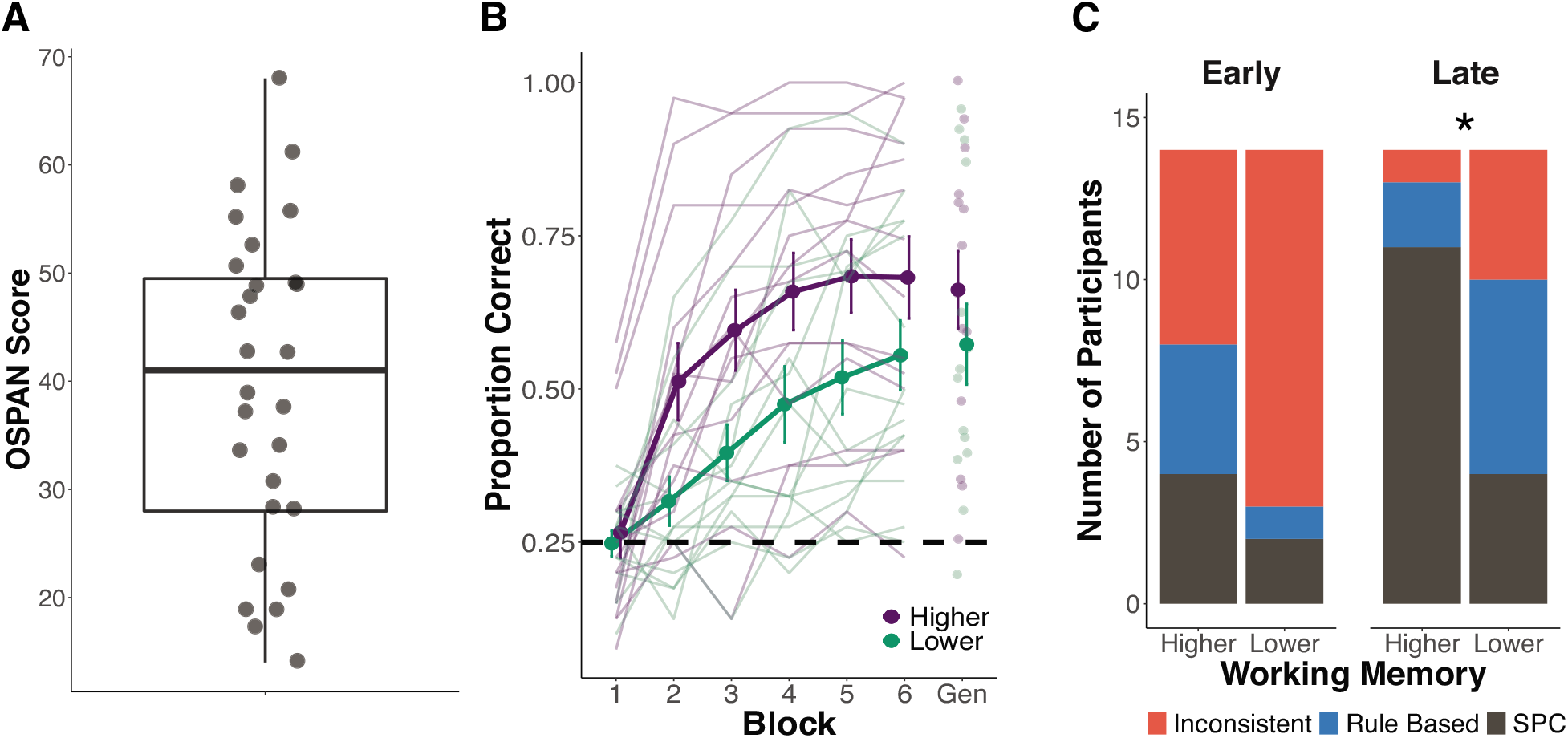
Experiment 2 behavioral results. A) Distribution of scores on the OSPAN task for all 28 participants. B) Categorization accuracy across training blocks and generalization block (abbreviated as “Gen”) for higher and lower working memory (WM) participants. Categorization accuracy is defined as the proportion of correct trials within each block. Average accuracy for each WM group is denoted by the darker lines and points. The lighter lines and points denote each individual participant’s accuracies. The dashed line reflects chance-level performance (0.25). C) Distribution of strategy usage among higher and lower WM participants in early and late training halves. There were no differences in strategy usage in early training between WM groups, but strategy usage significantly differed in late training between higher and lower WM participants (*p* = .034)

Participants used different strategies during category learning. We examined the extent to which categorization accuracy differed in early and late training between, Rule-Based strategy users (early: *M* = 0.475, *SD* = 0.232; late: *M* = 0.734, *SD* = 0.174), SPC strategy users (early: *M* = 0.594, *SD* = 0.174; late: *M* = 0.695, *SD* = 0.180), and Inconsistent strategy users (early: *M* = 0.382, *SD* = 0.121; late: *M* = 0.383, *SD* = 0.093), regardless of WM group membership. There was a significant effect of strategy type in early (*F*(2, 25) = 4.275, *p* = 0.025, η_p_^2^ = 0.255) and late training (*F*(2, 25) = 7.905, *p* = 0.002, η_p_^2^ = 0.387). Tukey-HSD post-hoc testing revealed that the mean categorization accuracy in early training of SPC strategy users was significantly greater than the categorization accuracy of Inconsistent strategy users. However, categorization of Rule-Based strategy users in early training did not significantly differ from Inconsistent or SPC strategy users. In late training, both Rule-Based and SPC strategy users had significantly greater accuracy than Inconsistent strategy users, but accuracy did not differ between Rule-Based and SPC strategy users.

There were no differences in strategy use among higher and lower WM participants in early training (Fisher’s Exact Test, *p* = 0.184). However, in late training, strategy usage differed among higher and lower WM participants (*p* = 0.034; Fig. 4C). Specifically, participants with lower WM used a mix of learning strategies, while participants with higher WM were more likely to use optimal procedural-based (i.e., SPC) learning strategies. Although strategy usage differed between higher and lower WM participants in late training, there were no differences in categorization accuracy within each learning strategy. For those using SPC learning strategies in late training, there were no significant differences in categorization accuracy between participants with higher WM (*Mdn* = 0.756) and participants with lower WM (*Mdn* = 0.554; *W* = 14.000, *p* = 0.327, *r* = 0.270). Additionally, there were no significant differences in late training for higher and lower WM participants using Rule-Based learning strategies (higher: *Mdn* = 0.825; lower: *Mdn* = 0.633; *W* = 2.500, *p* = 0.314, *r* = 0.415) nor Inconsistent learning strategies (higher: *Mdn* = 0.250; lower: *Mdn* = 0.308; *W* = 3.500, *p* = 0.468, *r* = 0.487).

#### Pupillometry

We first compared pupillary responses between sound trials and silence trials to confirm that pupillary responses on sound trials differed from silence trials as a proof of concept (Fig. 5A). Pupillary responses on sound trials were significantly different from silence trials on the intercept (β = 0.085, *t* = 14.511, *p* < .001), linear (β = 0.371, *t* = 102.825, *p* < 0.001), quadratic (β = −0.309, *t* = −85.513, *p* < 0.001), cubic (β = −0.038, *t* = −10.617, *p* < 0.001), and quartic (β = 0.093, *t* = 25.869, *p* < 0.001) terms. These results indicate that pupillary changes on sound trials were larger, increased more gradually over time, were stronger in curvature, and had significantly different secondary and tertiary inflection points. Silence trials were not included in the subsequent GCA model. Full model details are provided in Table 2.

**Figure 5.**
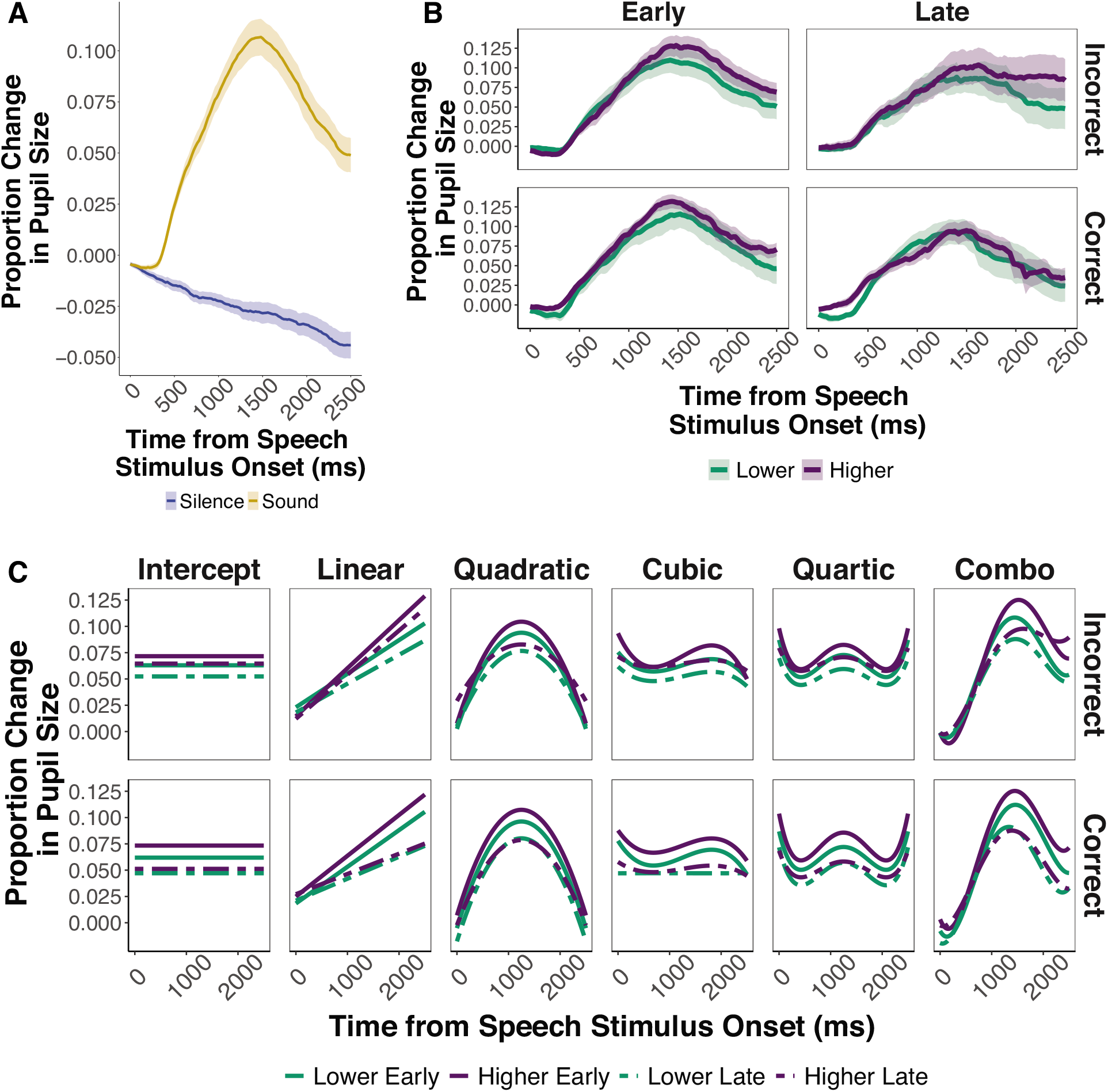
Pupillary responses (baseline normalized) to stimulus onset. A) Proportion change in pupil size for silence and sound trials. Shaded regions represent the standard error of the mean. B) Proportion change in pupil size as a function of working memory (WM) group for correct and incorrect trials in early and late training halves. Shaded regions represent the standard error of the mean. C) Decomposition by growth curve time terms of the proportion change in pupil size as a function of WM group and training half for participants with incorrect and correct trials. The combination growth curve (abbreviated as “Combo”) represents the sum of each individual time term component to model the pupillary response.

**Table 2.** Fixed effect estimates for model of pupillary responses to silence and sound trials from 0 ms to 2500 ms time-locked to stimulus onset (observations = 7056).

| **Fixed Effect** | **Estimate** | **SE** | ***t*** | ***p*** |
| --- | --- | --- | --- | --- |
| Intercept | -0.025 | 0.004 | -6.947 | < .001 *** |
| ot1 | -0.119 | 0.024 | -4.872 | < .001 *** |
| ot2 | 0.003 | 0.014 | 0.214 | .832 |
| ot3 | -0.013 | 0.009 | -1.377 | .179 |
| ot4 | -0.002 | 0.007 | -0.334 | .741 |
| Trial Type (Sound vs. Silence) | 0.085 | 0.006 | 14.511 | < .001 *** |
| ot1 x Trial Type | 0.371 | 0.004 | 102.825 | < .001 *** |
| ot2 x Trial Type | -0.309 | 0.004 | -85.513 | < .001 *** |
| ot3 x Trial Type | -0.038 | 0.004 | -10.617 | < .001 *** |
| ot4 x Trial Type | 0.093 | 0.004 | 25.869 | < .001 *** |

\* $p < .05$ , \*\* $p < .01$ , \*\*\* $p < .001$
Growth curve model: `modell <- lmer(Pupil ~ (ot1 + ot2 + ot3 + ot4) * TrialType + (ot1 + ot2 + ot3 + ot4 + TrialType|Subject), data = gca.data, control = lmerControl(optimizer = "bobyqa"), REML = FALSE)`

#### Effects of WM and trial accuracy in early training

Next, we examined pupillary responses in higher and lower WM participants for correct and incorrect trials in early training (see Table 3; Fig. 5B). For correct trials in lower WM participants in early training, we observed a significant, negative main effect on the quadratic term, indicating that correct trials had a steeper curvature than the pupillary response for incorrect trials in participants with lower WM in early training (Fig. 5C). For incorrect trials in participants with higher WM, no significant effects were observed on any of the terms, indicating that the pupillary response on incorrect trials in early training was similar in size, slope, curvature, and secondary and tertiary inflection points between higher and lower WM participants.

**Table 3.** Fixed effect estimates for model of pupillary responses from 0 ms to 2500 ms time-locked to speech stimulus onset to examine accuracy, half, and working memory (observations = 14,112).

| Fixed Effect | Estimate | SE | *t* | *p* |
| --- | --- | --- | --- | --- |
| Intercept | 0.062 | 0.008 | 7.375 | < .001 *** |
| ot1 | 0.273 | 0.061 | 4.432 | < .001 *** |
| ot2 | -0.328 | 0.029 | -11.313 | < .001 *** |
| ot3 | -0.063 | 0.0237 | -2.677 | .012 * |
| ot4 | 0.099 | 0.016 | 6.112 | < .001 *** |
| Accuracy (Correct vs. Incorrect) | -0.001 | 0.007 | -0.144 | .887 |
| ot1 x Accuracy | 0.024 | 0.012 | 1.937 | .053 |
| ot2 x Accuracy | -0.034 | 0.012 | -2.737 | .006 ** |
| ot3 x Accuracy | -0.015 | 0.012 | -1.241 | .215 |
| ot4 x Accuracy | 0.003 | 0.012 | 0.254 | .799 |
| Working Memory (Higher vs. Lower) | 0.010 | 0.012 | 0.840 | .408 |
| ot1 x Working Memory | 0.073 | 0.087 | 0.837 | .410 |
| ot2 x Working Memory | -0.008 | 0.041 | -0.185 | .855 |
| ot3 x Working Memory | -0.017 | 0.034 | -0.513 | .611 |
| ot4 x Working Memory | 0.016 | 0.023 | 0.680 | .501 |
| Accuracy x Working Memory | 0.003 | 0.010 | 0.266 | .792 |
| ot1 x Accuracy x Working Memory | -0.081 | 0.018 | -4.601 | < .001 *** |
| ot2 x Accuracy x Working Memory | 0.022 | 0.018 | 1.259 | .208 |
| ot3 x Accuracy x Working Memory | 0.048 | 0.018 | 2.746 | .006 ** |
| ot4 x Accuracy x Working Memory | 0.013 | 0.018 | 0.714 | .475 |
| Half (Late vs. Early) | -0.013 | 0.001 | -16.288 | < .001 *** |
| ot1 x Half | -0.075 | 0.009 | -8.611 | < .001 *** |
| ot2 x Half | 0.039 | 0.009 | 4.482 | < .001 *** |
| ot3 x Half | 0.043 | 0.009 | 4.875 | < .001 *** |
| ot4 x Half | -0.014 | 0.009 | -1.631 | .103 |
| Accuracy x Half | -0.004 | 0.001 | -2.699 | .007 ** |
| ot1 x Accuracy x Half | -0.078 | 0.018 | -4.423 | < .001 *** |
| ot2 x Accuracy x Half | -0.055 | 0.018 | -3.157 | .002 ** |
| ot3 x Accuracy x Half | 0.057 | 0.018 | 3.274 | .001 ** |
| ot4 x Accuracy x Half | 0.026 | 0.018 | 1.470 | .142 |
| Working Memory x Half | -0.002 | 0.001 | -1.697 | .090 |
| ot1 x Working Memory x Half | -0.020 | 0.012 | -1.609 | .108 |
| ot2 x Working Memory x Half | 0.067 | 0.012 | 5.418 | < .001 *** |
| ot3 x Working Memory x Half | 0.008 | 0.012 | 0.643 | .520 |
| ot4 x Working Memory x Half | -0.036 | 0.012 | -2.871 | .004 ** |
| Accuracy x Working Memory x Half | -0.011 | 0.002 | -4.926 | < .001 *** |
| ot1 x Accuracy x Working Memory x Half | -0.051 | 0.025 | -2.057 | .040 * |
| ot2 x Accuracy x Working Memory x Half | -0.025 | 0.025 | -1.022 | .307 |
| ot3 x Accuracy x Working Memory x Half | -0.089 | 0.025 | -3.600 | < .001 *** |
| ot4 x Accuracy x Working Memory x Half | -0.032 | 0.025 | -1.301 | .193 |

\**p* < .05, \*\**p* < .01, \*\*\**p* < .001
Growth curve model: *model2* <- *lmer*(*Pupil* ~ (*ot1* + *ot2* + *ot3* + *ot4*) \* *Accuracy* \*
*WorkingMemory* \* *Half* + (*ot1* + *ot2* + *ot3* + *ot4* + *Accuracy*|*Subject*), *data* = *gca.data*, *control* = *lmerControl*(*optimizer* = "bobyqa"), *REML* = *FALSE*)

In addition, significant two-way interactions between accuracy and WM were observed on the linear and cubic terms. For the linear term, the non-significant positive effect of correct trials on the linear term for lower WM was significant and negative for participants with higher WM. Similarly, for the cubic term, the significant effect of correct trials in early training in participants with higher WM was positive, but was non-significant and negative in participants with lower WM. These results indicate that the pupillary response on correct trials in early training for participants with higher WM were faster to dilate over time and had an earlier secondary inflection point than correct trials for participants with lower WM. These results suggest that pupillary responses on correct trials in early training may begin to reflect the switch to using procedural learning strategies in participants with higher WM.

#### Effects of WM between training halves and trial accuracy

Next, we examined the extent to which pupillary responses during accurate categorization were modulated across training between higher and lower WM participants (see Table 3; Fig. 5B). We observed significant main effects of training half on the intercept, linear, quadratic, and cubic terms, indicating that the pupillary response on incorrect trials during late training for participants with lower WM was smaller, increased at a slower rate over time, was shallower in curvature, and had an earlier secondary inflection point in comparison to incorrect trials in the early training half for participants with lower WM (Fig. 5C).

Additionally, working memory significantly interacted with training half on the quadratic and quartic terms, suggesting that participants with higher WM had a larger change in the curvature and tertiary inflection points of the pupillary response on incorrect trials from early to late training than lower WM participants. Simple effect analyses revealed that the pupillary response on incorrect trials for participants with higher WM became smaller in size (*t* = 6.380, *p* < .001), shallower in curvature (*t* = −11.841, *p* < .001), and the secondary (*t* = −5.375, *p* < .001) and tertiary (*t* = 3.762, *p* = .004) inflection points occurred earlier in time compared to early training. However, the rate of dilation over time did not differ (*t* > .05) from early to late training in participants with higher WM.

We also observed significant interactions between accuracy and training half on the intercept, linear, quadratic, and cubic terms for lower WM participants. Simple effect analyses revealed that on correct trials in late training, the pupillary response was smaller (*t* = 13.426, *p* < .001), increased less over time (*t* = 9.217, *p* < .001), had an earlier secondary inflection point (*t* = −5.762, *p* < .001), but no significant change in the curvature or tertiary inflection points (*t*s > .05), relative to correct trials in early training for lower WM participants. This same analysis also revealed that during late training for lower WM participants, the pupillary response for correct trials in late training increased at a slower rate over time (*t* = 4.318, *p* < .001), was steeper in curvature (*t* = 7.201, *p* < .001), and had an earlier secondary inflection point (*t* = −3.389, *p* = .016) than incorrect trials in late training. However, there were no significant differences in the overall size or tertiary inflection points (*t* > .05) between correct and incorrect trials in late training for lower WM participants. Collectively, these results indicate that the overall size, rate of dilation, and secondary inflection point of the pupillary response reduced from early to late training to a greater extent for correct trials in participants with lower WM participants, while incorrect trials had a greater reduction in the curvature of the pupillary response from early to late training compared to correct trials.

Finally, we observed significant negative three-way interactions between accuracy, WM, and training half on the intercept, linear, and cubic terms. Simple effect analyses revealed that on correct trials in late training for participants with higher WM, the overall pupillary response was significantly smaller (*t* = 20.048, *p* < .001), increased at a significantly slower rate over time (*t* = 12.883, *p* < .001), had a shallower curvature (*t* = −5.333, *p* < .001), and the tertiary inflection point occurred earlier in time (*t* = 4.286, *p* < .001) in late training relative to correct trials in early training, but the secondary inflection point of the pupil response did not significantly differ from early to late training (*t* > .05). This same analysis also revealed that pupillary responses on correct trials in late training for participants with higher WM increased at a significantly slower rate over time (*t* = 14.940, *p* < .001) and were steeper in curvature (*t* = 7.464, *p* < .001) compared to incorrect trials in late training, but the overall size, secondary, and tertiary inflection points did not differ between correct and incorrect trials in late training for participants with higher WM (*t*s > .05). Taken together, these significant three-way interactions indicate that participants with higher WM experienced a greater reduction in the pupillary response on correct trials from early to late training in the overall size and rate of dilation over time compared to participants with lower WM, while participants with lower WM experienced a greater reduction from early to late training on the secondary inflection point of the pupillary response on correct trials compared to participants with higher WM. Further, these significant three-way interactions demonstrate that in late training, participants with higher WM had a greater difference in the overall size between correct and incorrect trials than participants with lower WM. However, the difference in the secondary inflection point between correct and incorrect trials in late training was greater for participants with lower WM than those with higher WM.

Given that individuals with higher WM had greater overall learning accuracy in the speech category learning task, we also ran a third GCA identical to the accuracy, WM, and training half GCA reported above, except we included an additional random slope of trial count and subject to rule out the possibility that the quantity of correct and incorrect trials per WM group influenced our significant pupillary findings. The trial count variable consisted of the number of correct and incorrect trials per participant. This model did not change the results from the original model, demonstrating that our significant pupillary findings for correct and incorrect trials between participants with higher and lower WM during early and late training were not dependent upon the number of correct trials between WM groups.

Collectively, these results provide evidence that the effects of trial accuracy and WM on pupillary responses during speech category learning were modulated by training half. Specifically, pupillary response on trials that were categorized correctly reduced to a greater extent from early to late training in participants with higher WM compared to participants with lower WM. Participants with higher WM also had a larger difference in the pupillary response between trials categorized correctly and those categorized incorrectly in late training than participants with lower WM. These results demonstrate that pupillary responses may reflect greater use of optimal, procedural-based learning strategies for categorization by late training during speech category learning.

## General Discussion

We examined the extent to which individual differences in WM influence non-native speech category learning. In Experiment 1, we showed that adults with higher WM learned Mandarin tone categories better and faster than individuals with lower WM. In Experiment 2, we replicated these results in a smaller sample of participants who learned to categorize Mandarin tones while pupillometry was simultaneously recorded. Our neurobiologically-inspired computational models revealed that individuals with higher WM used more optimal, procedural-based strategies in late training, while those with lower WM perseverated with suboptimal, rule-based learning strategies. Pupillary responses revealed that participants with higher WM experienced a greater reduction in stimulus-evoked pupil size for correct trials by late training and showed larger differences between correct and incorrect trials in late training compared to participants with lower WM. Taken together, our findings indicate that higher WM promotes a faster switch to using optimal procedural-based learning strategies during non-native speech category learning. A substantial reduction in stimulus-evoked pupillary responses on correct trials during late training may further reflect a reduction in cognitive effort induced by the WM-independent, procedural-based strategy use.

The DLS framework posits that reflective and reflexive learning systems compete throughout learning (Chandrasekaran, Koslov, et al., 2014; Maddox & Chandrasekaran, 2014). Our results demonstrate that strategy use was similar between higher and lower WM individuals during early training, however, individuals with higher WM used more procedural-based learning strategies, mediated by the reflexive learning system, in late training compared to individuals with lower WM. While speech categories are argued to be optimally learned using the reflexive learning system and procedural-based learning strategies (Chandrasekaran, Koslov, et al., 2014; Yi et al., 2016), the reflective system that uses rule-based learning strategies and depends on WM, may still play a critical role in the learning process. Therefore, while WM may not have a well-defined role in the reflexive learning system itself, WM may facilitate the transition from the reflective learning system to the reflexive learning system for optimal speech category learning. In humans, there is a bias towards using rule-based learning strategies in early training, wherein successful learners switch to procedural-based learning strategies by late training, while less successful learners tend to continue using suboptimal rule-based learning strategies (Maddox & Chandrasekaran, 2014; Seger, 2008; Seger & Miller, 2010). Here, we observed that both WM groups did not primarily use a single learning strategy in early training, but individuals with higher WM were disproportionately inclined to use procedural-based learning strategies in late training. The established involvement of WM during rule-based learning (Decaro et al., 2008; Filoteo et al., 2010; Miles et al., 2014; Reetzke et al., 2016; Zeithamova & Maddox, 2006) may enable individuals with higher WM to better maintain speech stimulus-related information, which may in turn allow learners to extract and integrate key perceptual dimensions for accurate categorization.

Interestingly, these shifts in strategy usage were indexed by pupillary responses. Reductions in the stimulus-evoked pupillary responses across training reflected a reduction in cognitive effort, which corresponded with the shift to procedural-based learning strategies in individuals with higher WM. During early training, there were few stimulus-evoked pupillary differences between higher and lower WM individuals. By late training, differences were robust, wherein individuals with higher WM experienced a greater reduction in the overall size and rate of dilation on correct trials and had a greater difference between correct and incorrect trials in late training than individuals with lower WM. The reduction in the stimulus-evoked pupillary response across training on trials that were categorized correctly likely reflects the reduced cognitive effort (Kuchinsky et al., 2014; Morett et al., 2020; Winn et al., 2015) required for the reflexive learning system when categories are optimally learned using procedural-based learning strategies (Maddox & Chandrasekaran, 2014). Further, it is possible that the rate of the pupillary response over time (i.e., linear term) on correct trials may reflect the switch to using the reflexive learning system. In comparison to individuals with lower WM, individuals with higher WM had a faster rate of the pupillary response over time on correct trials in early training, experienced a greater reduction in the pupillary response over time on correct trials by late training, and had a significantly slower rate on correct trials relative to incorrect trials in late training. Taken together, individuals with higher WM experienced a greater change in their pupillary responses as a function of training that was associated with learning strategies.

Similar to prior neuroimaging findings that observed greater functional coupling between the left superior temporal gyrus and the striatum on incorrect responses during late training (Feng, Yi, et al., 2018), we observed greater pupillary differences between correct and incorrect trials for both higher and lower WM individuals in late training, in comparison to early training. Our findings demonstrate that incorrect trials had larger stimulus-evoked pupillary responses than correct trials for both WM groups in late training, but individuals with higher WM had a larger difference between correct and incorrect trials than individuals with lower WM. Interestingly, the increased coupling between the left superior temporal gyrus and the striatum on incorrect responses during late training in Feng et al. (2018) was observed during corrective feedback presentation. In the current study, we observed pupillary differences between correct and incorrect trials as early as the stimulus encoding period. While this suggests that both higher and lower WM individuals may have had greater attention to less established stimuli in late training, individuals with higher WM were better able to attend to the relevant dimensions of these less established stimuli to fine-tune their category representations in late training. These findings also demonstrated that a combination of corrective feedback and attention to relevant dimensions during stimulus encoding, modulated by individual differences in WM, drive speech category learning success.

Individual differences between individuals with higher and lower WM is purported to be caused by a deficit in LC-NE system functioning in individuals with lower WM (Unsworth & Robison, 2017a). One interpretation of these results is that a disruption in LC-NE functioning leads to attentional fluctuations in individuals with lower WM, such that individuals with higher WM were better able to attend to the speech stimuli and category-relevant dimensions. Thus, this enhanced attention in individuals with higher WM may facilitate the formation of novel category representations to a greater degree than in individuals with lower WM. Collectively, the differences in speech category learning performance, learning strategies, and pupillary responses observed between higher and lower WM individuals in the current set of experiments suggest that the LC-NE system is an important modulator of individual differences in WM during non-native speech category learning success. However, this study only included younger adult participants who were relatively within the normal ranges of WM. Further research should investigate nonnative speech category learning across the lifespan (i.e., in young children and older adult participants) and in populations with clinical deficits in WM.

Individuals with higher WM were better able to generalize to untrained talkers in Experiment 1, which may further indicate that individuals with higher WM not only attended to relevant dimensions but were also better able to incorporate corrective feedback to guide generalizable learning strategies. However, individuals with higher WM were not better able to generalize to untrained talkers in Experiment 2 in comparison to individuals with lower WM. The discrepancy in generalization performance between experiments may be driven by the nature of the speech category learning paradigm when paired with pupillometry. The duration of each trial component was extended in Experiment 2 to allow time for the slow-reacting pupillary response. This extended timing may have added an additional memory component to the task that was not present in Experiment 1, thus hampering the ability to generalize to untrained talkers.

In conclusion, we demonstrate that individual differences in WM facilitate non-native speech category learning in adults. Individuals with higher WM acquired non-native speech categories faster and with greater accuracies, relative to those with lower WM. Computational modeling and pupillometry revealed greater usage of procedural-based learning strategies and distinct stimulus-evoked pupillary signatures in individuals with higher WM as a function of training. We posit that individuals with lower WM may be more prone to attentional fluctuations, that may result in poorer encoding of key stimulus dimensions and suboptimal use of corrective feedback for error monitoring. The findings from the current set of experiments encourage further investigation of the role of individual differences in WM and the LC-NE system on non-native speech category learning success.

## Acknowledgements

This work used the Extreme Science and Engineering Discovery Environment (XSEDE, Towns et al., 2014), which is supported by NSF (ACI-1548562). Specifically, it used the Bridges system (Nystrom et al., 2015), which is supported by NSF (ACI-1445606), at the Pittsburgh Supercomputing Center.

